# Distinct neural computations scale the violation of expected reward and emotion in social transgressions

**DOI:** 10.1101/2024.04.29.591585

**Authors:** Ting Xu, Lei Zhang, Feng Zhou, Kun Fu, Xianyang Gan, Zhiyi Chen, Ran Zhang, Chunmei Lan, Lan Wang, Keith M Kendrick, Dezhong Yao, Benjamin Becker

## Abstract

Traditional decision-making models conceptualize humans as optimal learners aiming to maximize outcomes by leveraging reward prediction errors (PE). While violated emotional expectations (emotional PEs) have recently been formalized, the underlying neurofunctional basis and whether it differs from reward PEs remain unclear. Using a modified fMRI Ultimatum Game on n=43 participants we modelled reward and emotional PEs in response to unfair offers and subsequent punishment decisions. Computational modelling revealed distinct contributions of reward and emotional PEs to punishment decisions, with reward PE exerting a stronger impact. This process was neurofunctionally dissociable such that (1) reward engaged the dorsomedial prefrontal cortex while emotional experience recruited the anterior insula, (2) multivariate decoding accurately separated reward and emotional PEs. Predictive neural expressions of reward but not emotional PEs in fronto-insular systems predicted neurofunctional and behavioral punishment decisions. Overall, these findings suggest distinct neurocomputational processes underlie reward and emotional PEs which uniquely impact social decisions.

## INTRODUCTION

Scaling the discrepancy between actual and anticipated reward or punishment, generally referred to as prediction errors (PEs), critically guides social adaptive behavior which is essential for survival and personal development ^1^. Traditional learning and value-based decision-making models commonly posit that individuals acting as optimal learners strive to maximize their rewards while minimizing costs rely on reward PEs ^2,3^. This perspective is supported by convergent evidence from animal models that highlight the crucial role of dopamine reward PE signaling in driving approach learning towards rewarding stimuli ^4,5^. However, in humans the emotional reaction towards gains and losses may additionally impact decision making processes ^6^. Indeed, a positive PE (i.e., obtaining a better outcome than expected) reflects a pleasant surprise, elicits hedonic experiences and motivates the future pursuit of rewards, whereas a negative PE (i.e., obtaining a worse outcome than expected) evokes negative emotions such as disappointment and frustration, ultimately leading to avoidance ^4^. When making decisions, people engage in anticipation of the hedonic valence (pleasure or pain) associated with future outcomes. The accuracy of those prediction holds paramount importance, as an overestimation of pleasure pertaining to favorable outcomes can lead to risky choices, while an overestimation of the aversive experience for unfavorable outcomes may lead to avoidance and in turn missed opportunities ^11^. In an experimental context the corresponding decision process has been extensively examined using the classic social context-dependent Ultimatum Game (UG). In this economic exchange paradigm one player proposes a division of a sum of money and the other player (responder) can either accept or reject the offer (in which case neither the proposer nor responder receives any money). From an economic perspective the rational decision would be to accept even small offers to maximize reward, yet humans frequently reject offers that they consider unfair. The underlying decision making process has commonly been explained in terms of a negative reward PE signaling receiving the (lower) actual offer than expected and the behavior may serve to “punish” individuals who violate social fairness norms ^7^. However, humans do not solely build their models about the environment on reward computations and accumulating evidence indicates that humans establish complex mental models to accurately predict their own and others’ emotional experience ^8^, such that e.g. the anticipation of regret strongly impacts decision making ^9,10^.

While numerous studies have employed computational models to determine the behavioral and neural dynamics of classic reward PEs during social learning, initial studies indicate that anticipated emotions affect decisions. Within a reinforcement learning (RL) framework, the Rescorla-Wagner RL model appears suitable to explain social learning mechanisms ^11,12^. For instance, the learning rate at which people recalibrate their social expectations quantifies the extent to which PEs are integrated into the updating of reward values ^13^. In real-world social interactions, however, the basic RL model does not take into consideration the complex social contexts or associated emotional reactions. Sophisticated studies have begun to explore the impact of emotional experiences on RL-based learning in social contexts, demonstrating that empathy influences the prosocial learning rate ^14,15^ and acquiring knowledge about others’ emotions facilitates inferences regarding the informational value of social cues ^16^. A recent study by Heffner and colleagues utilized a modified version of the UG that required participants to report their anticipated and actual emotional state for (rather unfair) monetary offers and could demonstrate that emotional, rather than reward PEs are critical determinants for punishing unfair offers in an UG^17^. While these initial studies suggest a critical role of reward and emotional evaluations in social decisions, it remains unclear how the emotional and reward PEs are generated in the brain and uniquely shape decisions.

Although considerable evidence has outlined that a common ‘neural currency’ underlies evaluation of rewards ^18,19^, whether reward and emotional PEs are mediated by common or distinguishable neurofunctional computations has not been investigated. The striatum has been consistently reconciled as a center encoding reward PE during non-social learning, however in the context of social reward learning entailing additional social and emotional processes a broader brain network is recruited. When socially learning about other’s actions and outcomes, RL-like PEs with respect to expectations formed about how others viewed the self is revealed in the activity of anterior insula (aINS), anterior cingulate cortex (ACC) and orbitofrontal cortex (OFC)^20^. Additionally, the aINS has been proposed to mediate approach and avoidance in response to social affective stimuli ^21,22^, and critically contributes to avoidance of aversive emotional states ^23^. In social economic games such as UG where norm violations and aversive emotional states are intertwined, activation of the aINS has been associated with individuals’ willingness to reject unfair offers ^24^, possibly reflecting the involvement of the aINS in encoding both aversive and positive reward PEs ^25^. Moreover, numerous studies have explored the computational function of the ACC for encoding expectations and PEs related to others’ decisions during social RL learning ^26^, with prosocial PE signals in this region during learning to maximize monetary reward for others being correlated with the levels of empathy^27^. Encoding of social reward-related PEs by ACC neurons may further reflect the relevance of other individuals to one’s own emotional states, thereby eliciting changes in emotional arousal ^28^. Finally, the OFC is typically restricted to social context in a self-referenced framework such as encoding more the self-referenced type of reward PEs^29^. However, whether and how these systems process information about emotion and reward PEs as well as their distinct contributions to these two computational decision features remain to be explored.

Against this background we conducted a functional magnetic resonance imaging (fMRI) experiment (**Fig. 1a**) with a modified UG paradigm in N=43 healthy participants to examine the cognitive and neural mechanistic evidence of reward and emotional PEs underlying social value-based decision-making. In the modified UG paradigm, participants were required to predict how much reward they would get from the proposer and then predict how they feel on two emotional dimensions (valence and arousal) while again reporting their emotional experience when receiving the actual offer and finally deciding whether to accept or reject the proposer’s offer (**Fig. 1b**). When the offers are accepted, the monetary allocation will be made according to the offer, whereas when offers are rejected, neither the proposer nor responder receives any money (therefore often referred to as punishment decisions)^17^. On the behavioral level logistic mixed-effects regression models were employed to determine the predictive capacity of emotional and reward PEs to punishment decisions (**Fig. 1c**). On the neural level, we aimed to investigate: 1) the common and distinct neural systems that support the prediction and experience of reward and emotion during social decisions, 2) whether distinct multivariate neural patterns are sensitive to capture variations of emotional and reward PEs and the associated punishment decision using machine-learning based neural decoding approaches (given the higher precision of this approach to establish process specific neural signatures^30,31^), 3) whether the reward or emotional neurofunctional PE representation predict the neurofunctional decision to reject an offer and thus to characterize the different roles the PEs play in punishment decisions (**Fig. 1c**).

**Fig. 1.**
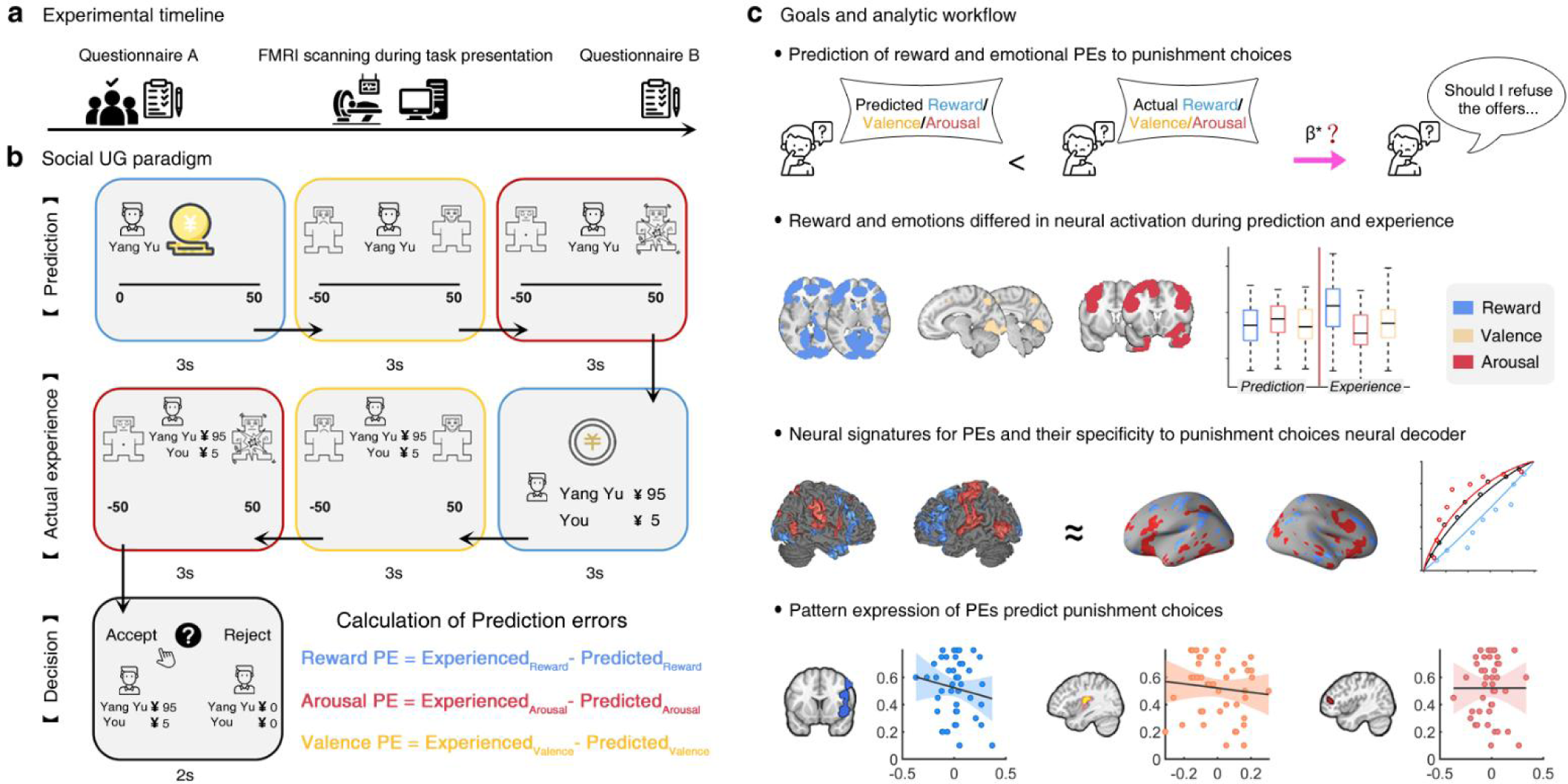
Experimental protocol, task design, and main goals and corresponding analytic workflow. **(a)** Experimental timeline. The questionnaire A and B include positive and negative affect scale, and state-trait anxiety inventory. **(b)** The modified Ultimatum Game task and PEs computation. Before each offer individuals predicted how much money they anticipated to receive from the proposers, and how they would feel in terms of valence and arousal when receiving the anticipated offer. After receiving the actual offers, individuals reported their current actual emotional experience in terms of arousal and valence and finally decided either to accept or to reject the offer. Main behavioral outcome where the computed reward and emotional PEs in terms of establishing trial-by-trial PEs: a reward PE (color coded in blue), an arousal PE (color coded in red) and a valence PE (color coded in yellow) scaling the difference between subjects’ prediction about the reward or emotion and their actual experience. **(c)** Major goals and analytic workflow of the current study. On the behavioral level, the predictive contribution of the reward and emotional PE to the decision to punish the proposer (i.e. rejecting the offers) was determined, modelling of the simultaneously acquired fMRI data aimed at: i) determining the univariate activation profiles of reward and emotional experience during the prediction and experience period to find separable neural underpinnings of reward and emotional, ii) training distinct multivariate neural patterns of emotional and reward PEs, as well as of punishment decisions to further indicate which neural pattern of PEs is specific to that of punishment decisions, iii) finally examining the links between the multivariate neural patterns for punishment decisions and all PEs to elucidate the neural pathway underlying the effects of reward and emotional PEs in social choices.

## Results

### Potential confounders

Both female and male participants were comparable with respect to the sociodemographic and pre-fMRI mood indices arguing against possible sex bias on the following results (all ps > 0.06, **Table 1**).

**Table 1.** Demographics and mood confounders

|  | Female (n = 23) | Male (n = 20) | T, p |
| --- | --- | --- | --- |
| Age | 21.57 ± 2.15 | 20.90 ± 2.07 | 1.03,0.31 |
| Body Mass Index,<br>kg/m<sup>2</sup> | 20.37 ± 2.31 | 21.56 ± 1.84 | -1.85,0.07 |
| PANAS-N | 16.70 ± 5.27 | 15.30 ± 3.92 | 0.97,0.34 |
| PANAS-P | 27.87 ± 5.96 | 29.75 ± 4.32 | -1.17,0.25 |
| TAI | 40.61 ± 9.03 | 37.25 ± 7.71 | 1.30,0.20 |
| SAI | 42.48 ± 9.43 | 39.55 ± 6.17 | 1.18,0.24 |

Values are presented as mean ± 1\*standard deviation. PANAS: Positive and negative affect scale, STAI-State and Trait anxiety inventory.

### Prediction of reward and emotional PEs to punishment decisions

In the realm of social interactions, complex social behaviors, such as the formation of alliances with peers, are theorized to be driven by the violation of expected outcomes including reward and emotions. We therefore tested the predictive role of reward and emotional PEs to the decisions about punishing a norm-violating proposer (i.e., rejecting the offers) via using a logistic mixed-effects regression model. Our results revealed that reward and emotional PEs significantly predicted punishment decision, such that participants showed higher punishment rates when experiencing less reward (β = -3.93 ± 0.46 (standard error), Z = -8.51, p < 0.001) and lower valence (β = -1.54 ± 0.29, Z = -5.27, p < 0.001) but higher arousal (β = 0.93 ± 0.20, Z = 4.71, p < 0.001) than anticipated (**Fig. 2a**). Moreover, we also employed β coefficient tests to compare the predictive power of reward and emotional PEs and found that the reward PE had a significantly stronger impact on motivating punishment choices than emotional PEs (reward PE vs valence PE, Z = -4.37, p < 0.001, reward PE vs arousal PE, Z = -8.89, p < 0.001). To explore the possible impact of sex on the observed findings the regression models were recomputed separately in male and female individuals. The results showed that all PEs robustly predicted punishment decisions (Female: reward PE, β = -4.03 ± 0.59, Z = -6.84, p < 0.001, valence PE, β = -1.71 ± 0.43, Z = -4.02, p < 0.001, arousal PE, β = 1.41 ± 0.38, Z = 3.68, p < 0.001; Male, reward PE, β = -3.93 ± 0.74, Z = -5.31, p < 0.001, valence PE, β = -1.47 ± 0.39, Z = -3.79, p < 0.001, arousal PE, β = 0.70 ± 0.18, Z = 3.83, p < 0.001; **Fig. 2a**) while the reward PE had a higher predictive capacity than emotional PEs (Female, reward PE vs valence PE, Z = -3.19, p < 0.001, reward PE vs arousal PE, Z = -7.48, p < 0.001; Male, reward PE vs valence PE, Z = -2.94, p < 0.01, reward PE vs arousal PE, Z = -5.54, p < 0.001). This result was consistent with the subsequent finding that female and male subjects exhibited a comparable reliance on all PEs to make punishment choices (reward PE × gender, β = 0.40 ± 0.79, Z = 0.50; valence PE × gender, β = 0.23 ± 0.53, Z = 0.43; arousal PE × gender, β = -0.44 ± 0.39, Z = -1.14; all ps > 0.25, **Fig. 2b**).

**Fig. 2.**
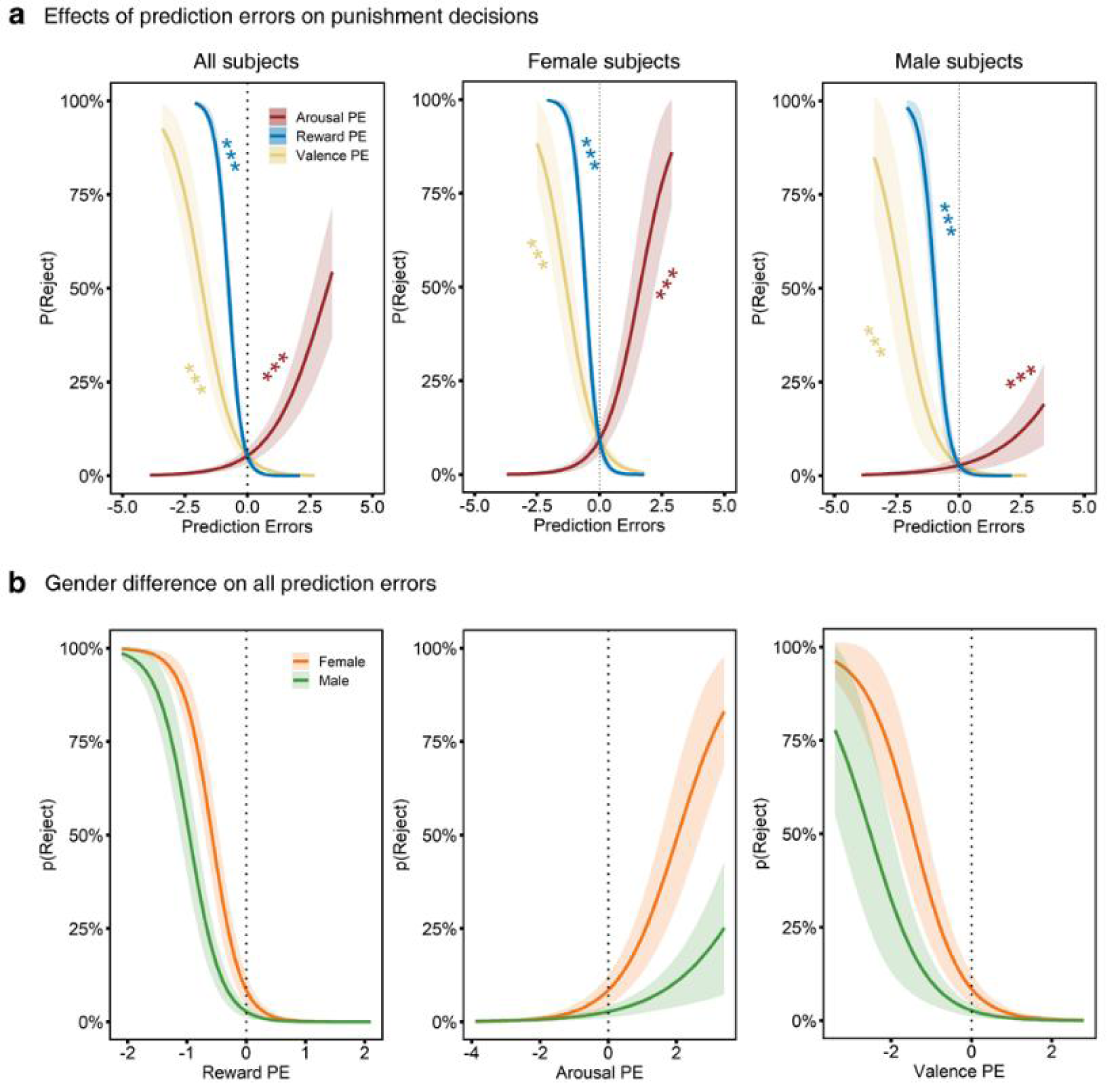
Predictive role of the different prediction errors for the decision to punish a social norm-violating proposer (reject an offer). **(a)** All PEs showed a significant predictive effects on punishment decisions such that participants showed higher punishment rates when they experienced less reward or lower valence but higher arousal than anticipated. **(b)** There was no significant gender difference on the reliance of all PEs to make punishment decisions. The lines reflect the probability of different choice pairs including rejecting versus accepting in the Ultimatum Game task, and the negative values represent negative PEs, suggesting less reward, less pleasantness, as well as less arousal than expected. Shaded areas reflect ± 1*standard errors. ***p < 0.001

### Neural activation for reward and emotion during prediction and experience stages

To further determine whether the anticipation and outcome evaluation on the reward and emotional level are supported by different brain systems we examined brain activation difference for reward and emotions separately for the prediction and experience stages of the experiment. While we did not observe that different regions were involved during predicting rewards and emotions, results suggested that the left dorsomedial prefrontal cortex (dmPFC, peak MNI coordinates, x/y/z = 10/48/46, F = 24.89, P_FWE-peak_ < 0.001, k = 566; **Fig. 3**) and bilateral aINS (left aINS, peak MNI coordinates, x/y/z = - 46/-2/4, F = 29.50, P_FWE-peak_ < 0.001, k = 267; right aINS, peak MNI coordinates, x/y/z = 52/6/10, F = 43.97, P_FWE-peakr_ < 0.001, k = 653; **Fig. 3**) were engaged differently during the reward and emotional evaluation stage. Examination of extracted parameter estimates (spherical masks, radius: 8 mm) revealed that the right dmPFC was activated stronger during experienced reward compared to emotions (**Fig. 3**), while the bilateral aINS was stronger engaged during valence and arousal experience as compared to reward experience (**Fig. 3**). Overall, these results indicate that the dmPFC encodes the experience of reward while the aINS encodes the actual emotional experiences in response to (unfair) offers.

**Fig. 3.**
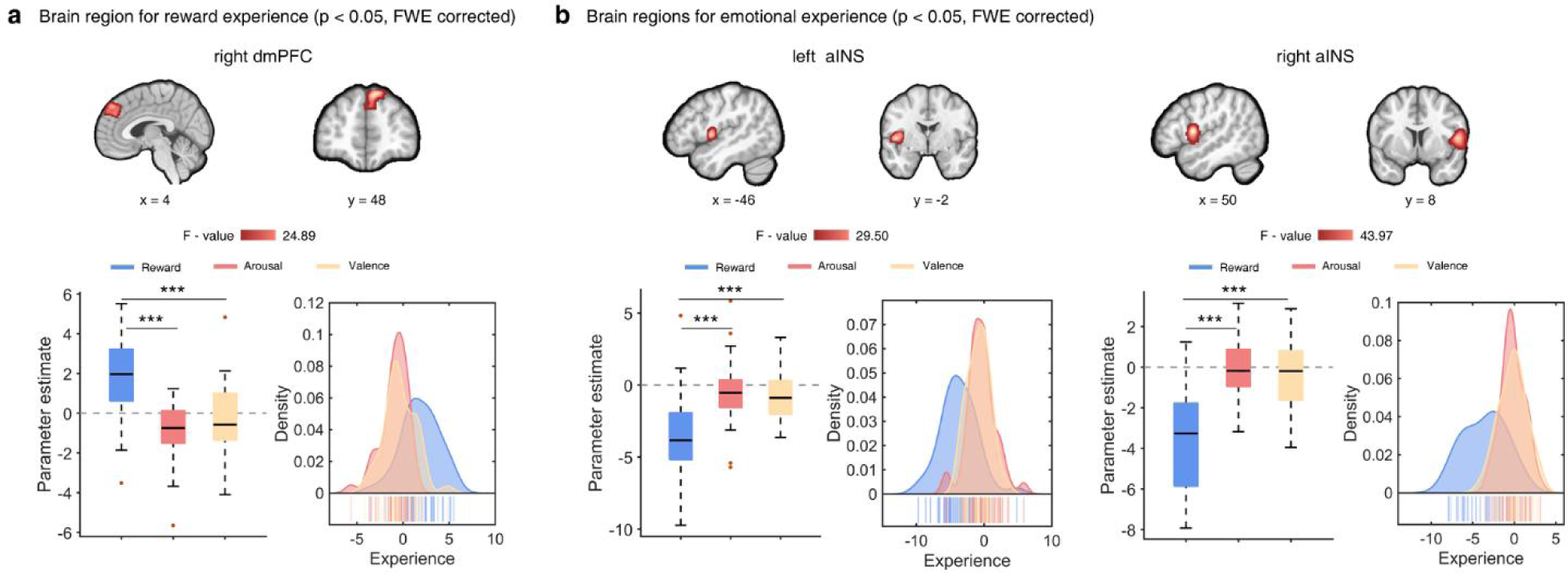
Brain regions processing the experience of reward and emotion, respectively. **(a)** The left dmPFC showed stronger engagement during reward relative to emotional experiences, while emotion processing regions such as the bilateral aINS showed higher activation for experienced emotions rather than the rewards **(b)**. For illustration purpose, parameter estimates were extracted from spherical (radius: 8 mm) regions of interest in the identified dosomedial prefrontal cortex (dmPFC) and anterior insula (aINS) regions. ***p < 0.001

### Neural signatures of reward and emotional PE

Given the high sensitivity of multivariate pattern analysis^32,33^ with respect to segregating cognitive and emotional processes we further developed a multivariate pattern classifier for reward and emotional PEs separated for punishment and accept decisions. Consistent with previous work, the decoding analysis revealed that multivariate predictive expression in the ventromedial prefrontal cortex, dorsolateral prefrontal cortex and dmPFC predicted the reward PE under accept decision (**Table S1**), while the multivariate expression in the right posterior insula (pINS), right ventrolateral prefrontal cortex (vlPFC), and left ACC was able to classify reward PE during punishment decision (accuracy, 0.72, sensitivity and specificity, 0.60, 0.84, separately, p < 0.001, **Fig. 4a**). Classification accuracy for emotional PEs that separated by punishment and accept decisions remained at chance level (all ps > 0.05).

**Fig. 4.**
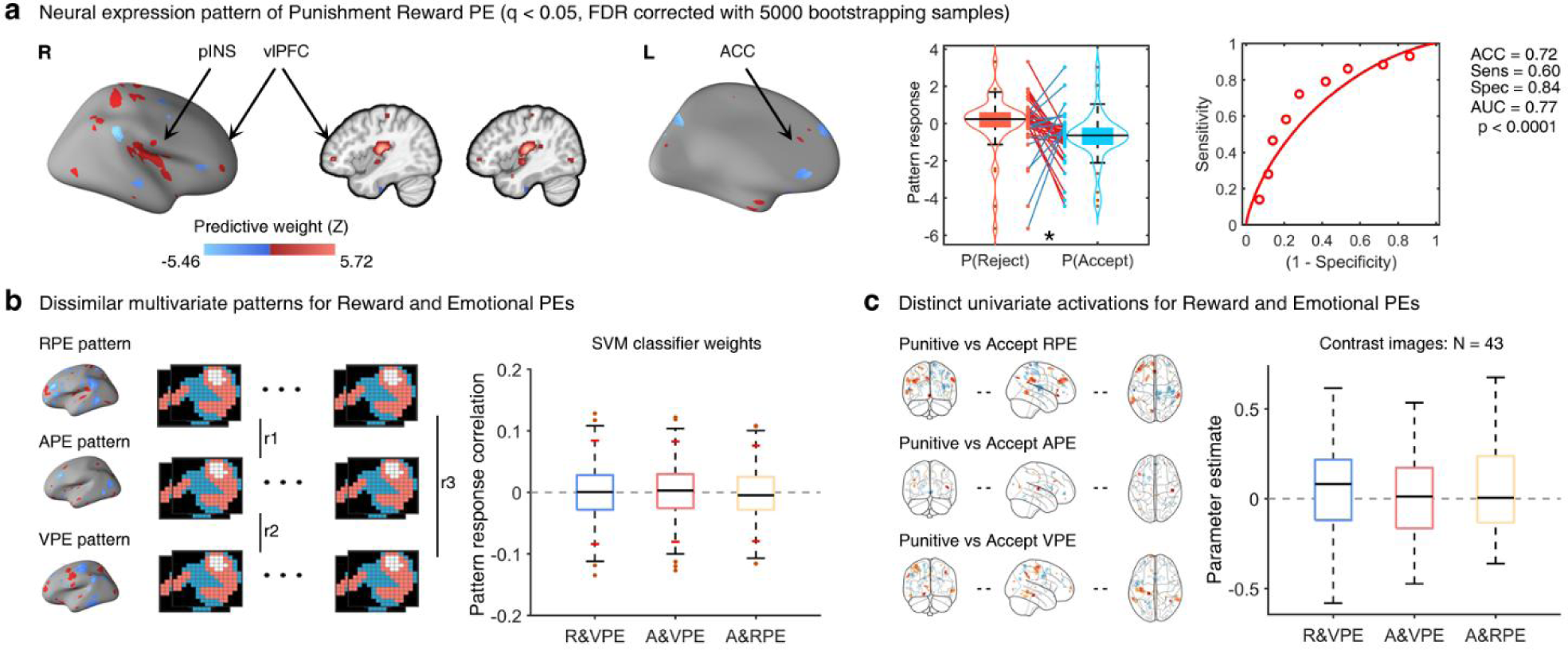
Multivariate neural expressions of reward and emotion prediction errors and their distinction. **(a)** Whole brain multivoxel pattern for differentiating punishment and accept reward PE with a bootstrapped samples (5000) which include the right pINS, right vlPFC and left ACC. The violin plot indicates that reward PE signature under punishment decision showed a higher response than that under accept decisions, t_(42)_ = 2.16, p = 0.04, two tailed, paired t-test, red line: high response in the reward PE under punishment decisions, blue line: high responses in the reward PE under accept decisions, and the forced-choice classification accuracy was 0.72, p < 0.001, two tailed, binomial test. **(b)** Bootstrapping test results for SVM classifier weight correlations. The short red lines reflect 95% confidence intervals obtained from bootstrap tests (500 samples). No regions showed significant correlations between SVM classifier weights. **(c)** The group-level correlations between activation of contrast images for punishment and accept-decision separating reward or emotional PEs. No regions showed significant average correlations between activations of contrast values across participants. ACC-accuracy, AUC-area under curve, Spec-specificity, Sens-sensitivity, *p < 0.05

To segregate the neural differences between reward and emotional PEs a pattern similarity analysis based on multivariate neural patterns of all PEs and a group-level correlation analysis between activations of contrast images for PEs were employed. We found no significant correlation between the whole brain multivariate expression weights for reward and emotional PEs (95% confidence intervals (CI) obtained from bootstrap tests (500 samples) all included zero) or contrast images for reward and emotional PEs for the specific decisions (all ps > 0.05, **Fig. 4b**). Together the findings indicate a sensitive neural pattern for reward PE in the frontal-insular circuit which was distinct from emotional PEs.

With the purpose of determining the (dis-)similarity between the reward and emotional PE representations, we fitted linear support vector regression (SVR) models to establish separate multivariate predictive signatures of reward and emotional PEs, and further examined their distinction. When using individual beta maps (one per PE level for each subject) as features to predict participants true PE value, we found overall correlations between predicted and actual reward and emotional PE values reached significance (0.31 < r < 0.48, all ps < 0.001) and the within-subjects prediction-outcome (i.e., 43×5=215 pairs) correlation coefficient was above 0.57 (**Fig. S2**). Moreover, classification accuracies for high versus low reward or emotional PE responses were higher than chance level in binomial tests (all ACC > 0.90, p < 0.001, **Fig. S2**). These findings demonstrate that during experiencing the actual rewards or emotions higher neural signature responses of the PEs were associated with stronger violations of the expected rewards or emotions, respectively. Consistent with previous classification results, the predictive neural signatures of reward and emotional PEs also showed spatial dissimilarity (reward & valence PEs, 95% CI, [-0.47, 0.43]; reward & arousal PEs, 95% CI, [-0.22, 0.22]; arousal & valence PEs, 95% CI, [-0.16, 0.14]).

### Univariate activation and multivariate expression pattern for Punishment versus Accept decisions

In line with previous studies systematically mapping neural activation related to social decisions^24,34^, we examined univariate activation for the differences between punishment and accept decisions and observed a fronto-insular network encompassing clusters located in the left dmPFC (peak MNI coordinates, x/y/z = -10/56/16, T = 6.13, P_FWE-cluster_ < 0.05, k = 996), bilateral ACC (peak MNI coordinates, left ACC: x/y/z = -10/48/14, T = 4.71, P_FWE-cluster_ < 0.05, k = 143, right ACC: x/y/z = 6/48/14, T = 4.05, P_FWE-cluster_ < 0.05, k = 76) and left aINS (peak MNI coordinates, x/y/z = -42/20/-10, T = 6.53, P_FWE-cluster_ < 0.05, k = 549, **Fig. 5a**) showing increased activation for punishment decisions. Further analyses using machine-learning based whole brain multivariate analyses^35^ via SVM classifier across individuals discriminated accept and reject decisions with high accuracy (0.78), high specificity (0.81) and high sensitivity (0.74). This neural pattern encompassed robust contributions of the left dmPFC, left aINS and left inferior frontal gyrus (bootstrapped 10,000 samples, FDR corrected, p < 0.05, **Fig. 5b**). In addition, given that reward and emotional PEs can directly predict the decisions to reject or accept the offers, we further explored whether the multivariate expression for punishment or accept decisions would be related to the PEs decoder pattern utilizing correlation analyses between the group-average reward or emotional PEs signatures responses separated by the decisions and the group-average decisions response. Results demonstrated – to a certain extent – the specificity of the reward PE pattern to predict the punishment decision expression (r = 0.29, p < 0.01, for emotional PEs all ps > 0.05).

**Fig. 5.**
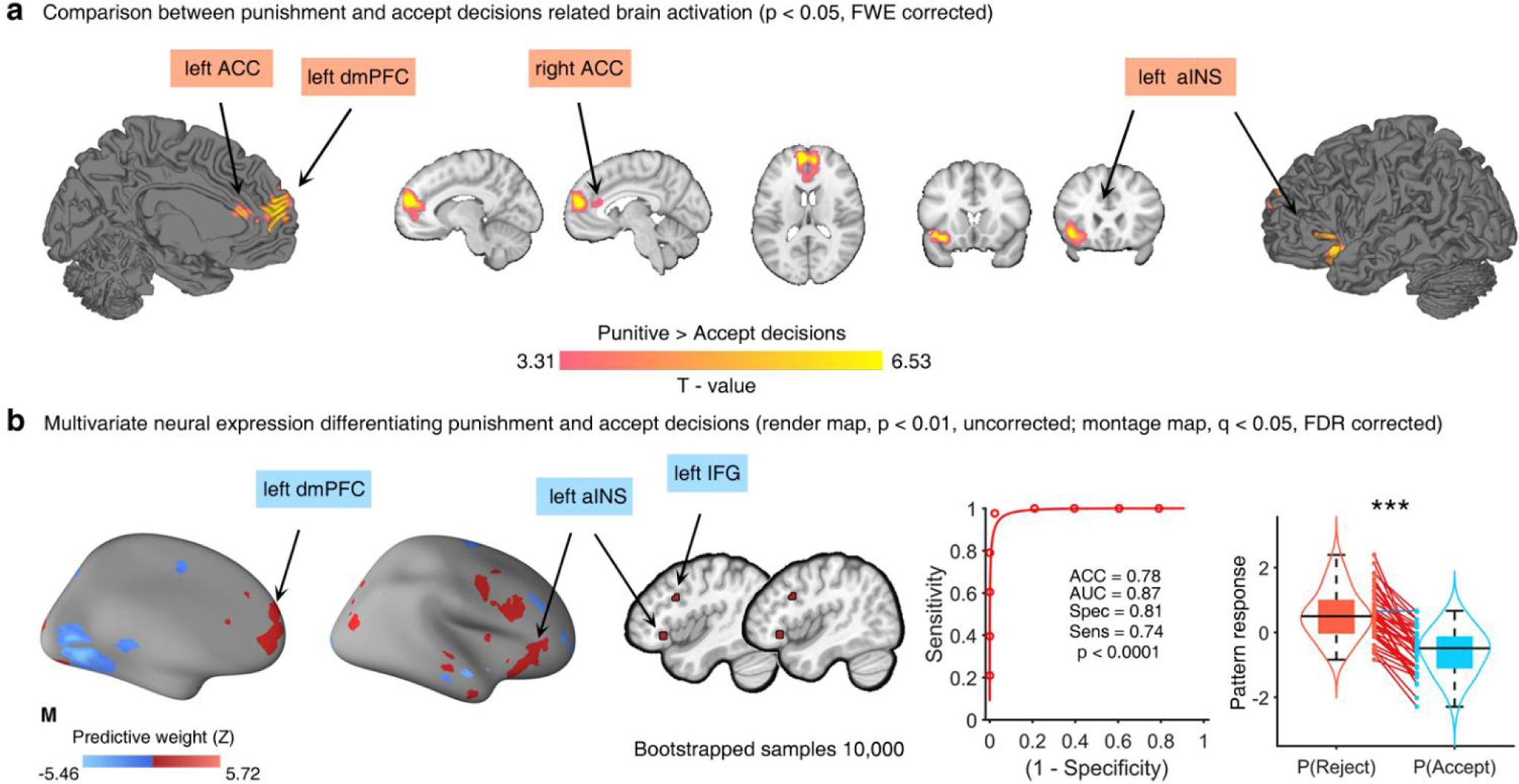
Univariate and multivariate patterns of punishment versus accept decisions. **(a)** Univariate activation for the difference between punishment and accept decisions showed that the bilateral ACC, left dmPFC and aINS were activated strongly for punishment choices (reject the offer). **(b)** Multivariate neural expression classifying the punishment and accept decisions included regions such as the left dmPFC, aINS and inferior frontal gyrus. The violin plot indicates that punishment decisions showed a higher level response compared to accept decisions, t_(42)_ = 10.85, p < 0.001, two tailed, paired t-test, red line: high response in the punishment decisions, blue line: high responses in the accept decisions, and the forced-choice classification accuracy was 0.78, p < 0.001, two tailed, binomial test. ACC-accuracy, AUC-area under curve, Spec-specificity, Sens-sensitivity, ***p < 0.001

### Pattern expressions within the reward PE decoder predict punishment decisions

Finally, to establish the functional association between neural signatures and behavior, we applied correlation analyses to investigate whether the neurofunctional PE signatures could predict punishment decisions on the behavioral level. We observed that pattern expressions for the reward PE under the punishment condition (r = 0.42, p < 0.01, **Fig. 6a**), but not for emotional PE (all ps > 0.10, **Fig. 6b**), encompassing a frontal-insular network were significantly positively associated with the decision to punish. A stronger pattern expression of the reward PE under the punishment condition predicted higher rates at rejecting unfair offers. Combined with our observation of a more sensitive signature for reward PE (punishment vs accept) and the close link of this decoder pattern with punishment decisions related multivariate neural expression, findings indicate that the reward PE (relative to emotional PEs) exerts a stronger influence on the decision to punish a proposer allocating unfair offers.

**Fig. 6.**
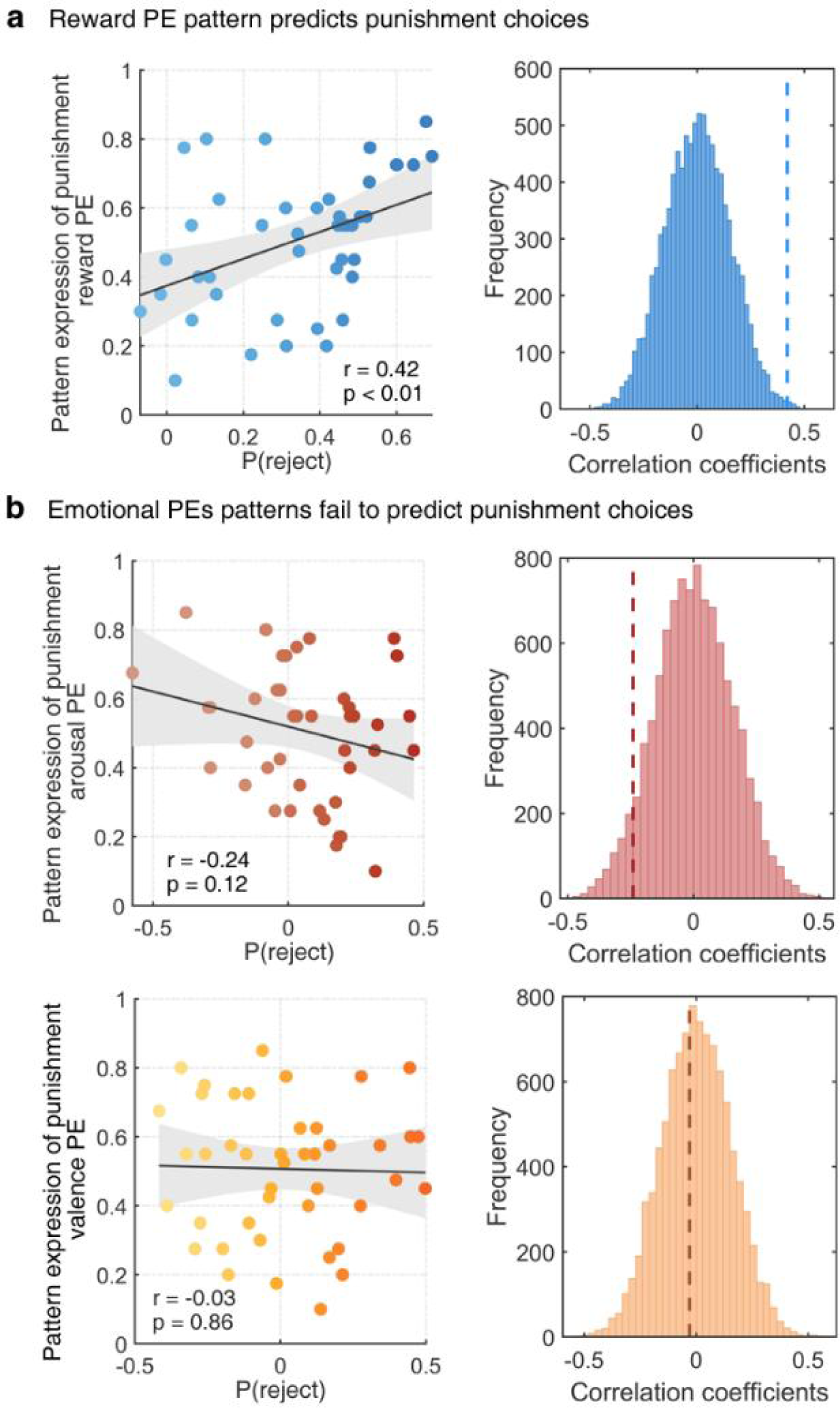
Exploratory correlation between brain and behavior. **(a)** The scatter plots reflect that stronger pattern expression within frontal-insular network representing punishment reward PE significantly correlate with increased number of punishment choices, while this association was diminished for emotional PEs **(b)**. The histograms show correlation coefficients from permutation tests, whereas the dashed lines represent the true correlation.

## Discussion

Traditional neurobiological decision-making models have determined the critical role of reward PEs signaling in fronto-striatal circuits in shaping choices and learning across contexts ^36^. However, the accompanying emotional response scaling the discrepancy between the expected and actual experiences as well as the underlying neurocomputational mechanism and whether they differ and interact with reward PEs has not been systematically determined. Here we combined a recently developed and modified UG paradigm which allows to quantify differences between experienced and predicted monetary reward and emotional experiences (along valence and arousal dimensions) in terms of reward and emotional PEs, respectively, with fMRI and multivariate predictive neurofunctional decoding to determine common and distinct influences of reward and emotional PEs on the decision to punish an unfair offer and the underlying neurocomputational representations. Our behavioral results confirmed that both reward and emotional PEs could significantly predict punishment decisions, with participants punishing at higher rates when experiencing less reward or pleasantness or more arousal than expected. However, the reward PE exerted a comparably stronger impact on motivating punishment choices compared to both emotional PEs. On the neural level, reward and emotional processes exhibited distinct neural representations across univariate and multivariate analyses. The experience of reward engaged the left dmPFC whereas the bilateral aINS was engaged during emotional experiences. Moreover, reward and emotional PEs were encoded in distinguishable brain-wide neural patterns. During the decision stage a fronto-insular network including the bilateral ACC, left aINS, left dmPFC and IFG increased activation and neural expression for punishment decisions, while the multivariate signatures of these regions closely resembled the distributed reward, yet not emotional PE signatures. In support of this an exploratory correlation analysis further demonstrated that a higher fronto-insular pattern expression under punishment reward PE predicted the subsequent punishment decision following unfair offers. Taken together, these findings shed lights on the computational and neural mechanism that distinguish emotional and rewards evaluation and how violations of expected rewards and emotions determine social decision making.

Rewards serve as an essential motivational driver that shapes adaptive decisions to optimize reward outcomes and to avoid negative consequences ^37^. Classical models of learning and decision-making have focused solely on reward and conceptualized individuals as optimal learners striving for maximizing expected rewards ^38–40^. Despite the normative appeal of those models in describing how humans evaluate and decide based on the difference between expected and actual reward outcomes, these models do not provide a clear psychological mechanism to account for the accompanying affective processes (e.g., emotion) and their influence on decision making. Punishment and uncooperative decisions are driven by a diverse array of negative emotions, including sadness and disappointed ^41^. In relation to reward computations a positive reward PE signaling that the experienced reward exceeds expectations can evoke pleasant feelings while a negative reward PE may induce strong negative emotions such as disappointment or anger ^4^. Utilizing computational modeling recent studies found that momentary emotions are not only explained by reward outcomes but rather by the combined influence of reward expectations and PEs that depend on these expectations ^39,42^. Together this suggests that reward PEs shape not only future decisions but also the momentary emotional state which in turn may impact the decisions. In line with this hypothesis a recent study demonstrated that violations of expected emotions (in particular valence) linked with expected and experienced rewards predict punishment choices in response to unfair offers stronger than reward PEs ^17^. Partly consistent with this work, our results suggest that emotional and reward PEs are intertwined to guide social decisions, such that both violations of reward and emotional predictions lead to punishment choices towards unfair offers.

On the neural level, we initially examined whether the emotional and reward processes are localized in separable neural systems and found that univariate activation in the dmPFC or aINS characterized experienced reward or emotion, respectively. Numerous previous neuroimaging studies have suggested an important role of the dmPFC in metacognitive effort cost valuation during reward decisions ^43,44^. However, this view has been recently challenged by findings indicating that the dmPFC is a crucial site for deploying learnt reward values in action selection particularly during social inference ^45,46^. Supporting this role of the dmPFC, a recent rodent study reported dmPFC neuronal activity accurately predicting reward availability and initiation of conditioned reward seeking after cue-reward learning ^47^. In contrast, the aINS exhibited a stronger engagement during the report of actual emotional experience upon receiving the offer. In the context of unfair monetary treatments, people would generate negative emotions especially when they obtained less reward amount than expected. This may increase the activation of aINS which plays an important role in the evaluation and experience of aversive experiences, such as pain, disgust ^48^, introception and approach-avoidance decisions in social contexts ^22,49^ and witnessing unfair transactions or unmoral acts ^24,50^.

Consistent with results from univariate analyses, multivariate predictive modelling also revealed distinct distributed neural representations of emotional and reward PEs. However, we did not obtain a multivariate decoder that could robustly and sensitively capture variation of emotional PEs. Despite that, our recent study utilizing the data from this modified UG paradigm have determined that unfair offers indeed evoke a strong aversive emotional response within subcortical regions for avoidance responses (amygdala, PAG, thalamus, putamen) and cortical systems involved in emotional appraisal such as the insula, dorsal ACC and lateral frontal regions ^30^. This may provide some inspiration for understanding the possible neural pathways underlie the emotional PEs and could further support spatial dissimilarity between emotional and reward PEs, since the reward PE was implicated in a concentrated frontal-insular network encompassing the left ACC, right vlPFC and pINS. These findings align with previous studies that negative reward PE followed by unfair offers are embedded in the typical brain regions engaged in generating PE-like signaling for unexpected reward outcomes (ACC)^51^ and anticipation of uncertain punishments or rewards with the same neural codes (vlPFC) ^52^. Our results, however, extend more broadly to indicate that negative reward PE computed based on the unfair offers is also represented in the region for interoceptive processes ^53^ and exerting regulatory top-down control over reward-related behaviors via its projections to the nucleus accumbens ^23^.

We further determined whether the fronto-insular network encompassing the dmPFC, aINS, ACC and IFG represents or predicts punishment decisions via employing univariate and multivariate analytic approaches. The dmPFC and aINS exhibited the most stable predictive weights across univariate and multivariate analyses. These findings in line with the previously described role of the dmPFC in personal moral decisions via serving as a conflict monitor for inequality of economic offers ^54,55^ and self-disadvantageous unfairness ^56^. During human cooperation, the dmPFC is important for modulating outcome value signals for oneself and others to guide behavior appropriate to the local social context ^57^. Similar to the functions of dmPFC in social decisions, the aINS holds independent neural populations responding specifically to aversive states in a disadvantageous-inequality processing ^48,58^ and showing heightened activity when rejecting unfair offers ^24,59^. In particular, the functional integration between ACC and aINS is enhanced when individuals make dishonest decisions for a monetary reward, as it requires high need for emotional processing and conflicts detection ^60^ which can also be generated during evaluating outcomes of self and proposers in our task. Significant activation of aINS and IFG can be simultaneously observed when asking subjects to reappraise the proposers’ money dividing intentions as more negative and is linked with a greater number of unfair offers rejected ^61^. However currently the specific contributions of the separate systems in terms of punishment decisions during norm violations remain unclear. Our findings may bridge this gap by determining which PE representation significantly predict punishment choices. Within this context our predictive models demonstrated high specificity of fronto-insular neural predictive weights for the reward but not the emotional PEs to the multivariate response of punishment decisions. Moreover, we revealed that aINS and ACC generally engaged in unfairness perception and rejection of unfair offers given that they demonstrated increased pattern expression of punishment reward PE to significantly predict behavioral punishment choices. Overall, these findings underscore the critical role of reward PE coding in the fronto-insula system on decisions in response to unfairness.

Our study extends traditional learning and economic theories ^38,40^ by highlighting the crucial contribution of both violations of expected emotion and rewards to the decisions. The prevailing theories posit that humans make decisions via assigning values to prospective gains or loss ^40^, and have only implicated the activation within frontal-striatal circuits in representing value of anticipated or received rewards ^36^. We updated this view by demonstrating that the neurofunctional computation of reward is accompanied by simultaneous computations of emotional states and further showed that these processes are supported by distinguishable brain systems, such that the dmPFC encoded the receipt of the actual reward whereas the aINS encoded experienced emotions. Utilizing a more rigorous machine-learning based neural decoding method, the multivariate neural patterns between reward and emotional PEs were also found dissimilar. In general, our findings extend influential theories of decisions which mainly focused on reward effects on choice behaviors and emphasize the importance of considering emotions in economic models, especially how these factors are dissociable from neural pathways during social decisions.

It is worth highlighting potential limitations of the present study. Although we demonstrated consistent behavioral results via the use of multiple logistic regression models ^17^, the dynamic changes of subjects’ reliance on those PEs to make final decisions are not clear. This concern could be well resolved if future studies employed computational modeling to track the trial-wise updates of PEs and the relevant decision variations. Additionally, no direct modulation of the underlying neural processes was included and future studies may consider employing pharmacological approaches to regulate associated signaling systems such as the dopamine ^62^, oxytocin ^63^ or angiotensin ^64,65^ and to determine whether separable signaling systems underlie emotional and reward PEs.

In conclusion, our study supports the dissociable contribution of emotion and reward evaluations to social decisions by identifying distinct brain regions engaged in experienced emotions and rewards, and proving dissimilar multivariate neural patterns between PEs generated from those experiences. Despite the contribution of both emotional and reward PEs to social decision making, the latter could strongly guide social decisions given that the frontal-insular predictive neural expression of reward PE specifically correlates with multivariate pattern of punishment decisions and predict behavioral punishment choices. As such, our findings pave the way towards more precise understanding about how emotions and rewards distinctively represented in the brain to affect social decisions, and may have further implications on clinical disorders with deficient reward and emotion processing in complex social contexts.

## Methods

### Participants

The study was approved by the Research Ethics Committee of the University of Electronic Science and Technology of China (1061422101024711) and adhered to the latest revision of the Declaration of Helsinki. We employed an fMRI experimental design with N = 50 (25 Female, 25 Male) individuals. A total of 7 subjects were excluded due to fMRI data acquisition failure (fMRI technique issue and withdraw during the experiment, n = 3) and no punishment decisions (not rejecting the unfair offer, n = 4), leading to a final sample of N = 43 (23 Female, Mean ± SD, age = 21.57 ± 2.15 years; 20 Male, age = 20.90 ± 2.07 years) included into main analyses.

Exclusion criteria for enrollment were: (1) an excessive head movement (>2 mm translation or 2° rotation), (2) a current or a history of psychiatric, neurological, or other medical disorders, (3) current or regular use of psychotropic substances including nicotine, (4) a body mass index < 18 or > 24.9, (5) visual or motor impairments, and (6) contraindications for MRI.

### Experimental paradigm

An adapted and validated reward-emotion UG paradigm was employed ^17^(**Fig. 1**). Similar to the classical UG task participants were responders who received an unfair monetary offer (split) from a proposer, and then decided whether to accept or reject the offer in the modified version ^17^. However, more specifically, participant were instructed to give the following responses before each offer: 1) how much money (within a range of ¥2.5 to ¥50, 20 offers; ¥, Chinese Yuan) they would expect to receive from the proposer; and 2) rating the emotions that they expected to feel based on their anticipated monetary split along the two dimensions, valence (i.e., positive vs negative emotion) and arousal (i.e., intensity/strength of emotion). Following the offer all individuals reported: 1) their actual emotional experience upon receiving the offer; and 2) decided whether to accept or reject the proposer’s offers. A total of 40 task trials – dispersed across two fMRI runs with 20 trials each (2 trials per offer, trial mean duration 3s) were presented. Each task trial began with a fixation cross presented for a jitter interval of 500ms followed by the presentation of expected money and anticipated emotion, actual offer and experienced emotions, as well as the decisions periods. The offers followed a uniform distribution allowing each participant to respond to the full range of fair (¥50 vs ¥50) to unfair offers (¥97.5 vs ¥2.5), whereas the order of the offers was randomized.

### MRI acquisition, preprocessing and first level analysis

MRI data were acquired on a 3T GE Discovery MR system (General Electric Medical System, Milwaukee, WI, USA) and preprocessed using standard workflow in SPM 12 (Statistical Parametric Mapping; http://www.fil.ion.ucl.ac.uk/spm/; Welcome Trust Centre for Neuroimaging) (see **Supplemental Methods**).

To explore the brain regions involved in anticipated and experienced reward/emotion, and punishment/accept decisions, we firstly establish a general linear model (GLM) incorporating separate onsets of prediction or experience of reward, arousal or valence, respectively, as well as the decision period, while the six head motion parameters were included as covariates. To further determine the specific neural expressions for reward and emotional PEs in the context of accept or punishment decisions a separate parametric GLM was modelled including the experienced rewards, arousal and valence events for punishment and accept decisions as regressors, each modulated by the respective trial-wise PE.

### Behavioral analyses

With the purpose of evaluating predictive contribution of all PEs to punishment decisions simultaneously we employed a logistic mixed-effects regression model with three PEs as independent variables and punishment choices as the dependent variable via the lme4 R package (https://cran.r-project.org/). The computation of PEs was based on the disparity between the actual experience at the time of offers and prior expectation. It should be noted that a zero value of PEs indicated instances where participants’ actual experiences corresponded with their expectations, resulting in the absence of errors. As such all PEs were firstly scaled (i.e., PE/ sqrt(sum(PE^2)/length(PE)-1)) but not mean-centered before being included to the regression model to ensure that β coefficients could be comparable. To account for multicollinearity in regression models we first estimated the variance inflation factor (VIF) ^66^ (similar approach see Xu et al., 2020, Li et al., 2019^67,68^) with results arguing against problematic collinearity (VIF_Reward_PE_ = 1.52, VIF_Arousal_PE_ = 1.60, VIF_Valence_PE_ = 1.21). We also performed separate regression analyses in female and male subjects to avoid possible gender bias on PEs’ predictive role to punishment decisions.

To further determine the efficiency of the current paradigm and to further control gender effects, we scaled the difference between proposer’s and responder’s reward amounts in terms of unfairness (e.g., ¥92.5 - ¥2.5 = ¥90) and included it with gender as predictors for punishments choices in the same regression model. Unfairness significantly predicted the punishment decisions with the effect being equal in female and male subjects (Gender × Unfairness, β = 0.03, z = 0.05, p = 0.96, **Fig. S1**). This confirms a close association between the unfairness manipulation and the punishment decision across both genders.

### Univariate voxel-wise analyses

Considering the pivotal role of reward and emotions in guiding decisions, we aim to identify whether the neural activations of reward and emotion were distinct during prediction and experience moments. Given the strengths of univariate voxel-wise analyses in terms of determining the spatial localization of mental processes^69^, we initially employed this method to examine brain regions that were differentially engaged for reward and emotions (i.e., arousal and valence) during prediction and experience periods, respectively, via including the first level contrasts (i.e., Prediction_reward_, Prediction_arousal_, Prediction_valence_; Experience_reward_, Experience_arousal_, Experience_valence_) into two separate univariate voxel vise one-way ANOVAs. All resulting maps from the whole brain analyses were thresholded at the peak level of family wise error (FWE) correction at p < 0.05. Furthermore, to assess the neural activations differences between accepting and punishing decisions, we utilized the univariate voxel wise one sample t test on the first level contrast (i.e., Punishment decision > Accept decision) with the resulting t-value maps being thresholded at the cluster level of FWE correction at p < 0.05 (initial cluster threshold, p < 0.001, uncorrected; see recommendations in Slotnick, 2017^70^).

### Multivariate voxel pattern analyses

Compared to the traditional univariate analyses, the machine-learning based multivariate pattern analyses can provide more comprehensive and precise neural representations of cognitive and mental processes^30,31^. Therefore we utilized a linear support vector machine (C = 1, linear kernel) implemented in Canlabcore tool (https://github.com/canlab/CanlabCore) with a leave-one-out cross validation procedure to get differentiate neural patterns of reward and emotional PEs separated by punishment and accept decisions. Moreover, we also assessed whether the neural expressions of reward and emotional PEs were similar via conducting pattern similarity analyses between multivariate neural patterns of all PES and calculating group-level correlations between activations of contrast images for PEs based on bootstrap tests with 500 iterations.

To build liner neural expression maps predictive of the levels of reward and emotional PEs, we established exploratory GLM models to get different beta map for each reward and emotional PE level. The 20 reward and emotional PEs were sorted with descend sequence and were then divided into 5 levels (i.e., 5,4,3,2,1) in each run for each subject. Next, we employed support vector regression analyses using a linear kernel (C= 1) (in line with our previous works^30,31^) implemented in the Spider toolbox (http://people.kyb.tuebingen.mpg.de/spider) with individual beta maps (one per PE level for each subject) as features to predict participants true PE value. To facilitate a robust determination of the predictive accuracy of the neurofunctional signature we employed various metrics including correlation and forced-choice classification accuracy. We used overall (between- and within-subjects; 43 × 5 = 215 pairs in total) and within-subject (5 pairs per subject) Pearson correlations between the cross-validated predictions and the actual PEs to indicate the effect sizes. We also assessed classification accuracy of the signatures of PEs using a forced-choice test, where signature responses were compared for two conditions tested within the same individual, and the higher was indicated large violations of predicted emotions or rewards (see **Supplemental Methods**).

### Exploratory correlation analysis

The behavioral and fMRI results suggested a pronounced predictive role of reward PE to punishment decisions. We thus conducted an exploratory correlation analyses with permutation tests (10,000 permutations) to identify whether multivariate pattern expressions for reward PE generated before punishment decisions could predict behavioral punishment choices. For the sake of completeness and increasing transparency we also reported the correlations between punishment decisions and the multivariate pattern expressions for emotional PEs signaling.

## Acknowledgement

This work was supported by the China MOST2030 Brain Project (Grant No. 2022ZD0208500), National Natural Science Foundation of China (Grants No. 32250610208, 82271583), and National Key Research and Development Program of China (Grant No. 2018YFA0701400) and a start-up grant from The University of Hong Kong. Disclaimer: Any opinions, findings, conclusions or recommendations expressed in this publication do not reflect the views of the Government of the Hong Kong Special Administrative Region or the Innovation and Technology Commission. We truly thank all colleagues for the important discussions that helped us to improve the manuscript and the help during the fMRI data acquisition, and all volunteers who participated in our study.

## Author contributions

TX and BB designed the study. TX, KF, LW, RZ, ML conducted the experiment and collected the data. TX performed the data analysis with important advice provided by FZ, LZ, YG and YC. TX and BB wrote the manuscript draft which was further critically revised by LZ, YC, KK and ZY.

## Data and code availability

The multivariate brain patterns and source behavioral data for figures will be shared upon publication through a Figshare repository. The code for generating the figures and main analyses will be shared upon publication through a GitHub repository.

## References

1 Cohen, J. Y., Haesler, S., Vong, L., Lowell, B. B. & Uchida, N. Neuron-type-specific signals for reward and punishment in the ventral tegmental area. nature 482, 85–88 (2012).

2 Sambrook, T. D. & Goslin, J. A neural reward prediction error revealed by a meta-analysis of ERPs using great grand averages. Psychological bulletin 141, 213 (2015).

3 Tymula, A. et al. Dynamic prospect theory: Two core decision theories coexist in the gambling behavior of monkeys and humans. Science Advances 9, eade7972 (2023).

4 Schultz, W. Reward prediction error. Current Biology 27, R369–R371 (2017).

5 Schultz, W. Dopamine reward prediction-error signalling: a two-component response. Nature reviews neuroscience 17, 183–195 (2016).

6 Paulus, M. P. & Angela, J. Y. Emotion and decision-making: affect-driven belief systems in anxiety and depression. Trends in cognitive sciences 16, 476–483 (2012).

7 Van’t Wout, M., Kahn, R. S., Sanfey, A. G. & Aleman, A. Affective state and decision-making in the ultimatum game. Experimental brain research 169, 564–568 (2006).

8 Thornton, M. A. & Tamir, D. I. Mental models accurately predict emotion transitions. Proceedings of the National Academy of Sciences 114, 5982–5987 (2017).

9 Coricelli, G. et al. Regret and its avoidance: a neuroimaging study of choice behavior. Nature neuroscience 8, 1255–1262 (2005).

10 Mellers, B. A. & McGraw, A. P. Anticipated emotions as guides to choice. Current directions in psychological science 10, 210–214 (2001).

11 FeldmanHall, O. & Nassar, M. R. The computational challenge of social learning. Trends in Cognitive Sciences 25, 1045–1057 (2021).

12 Frolichs, K. M., Rosenblau, G. & Korn, C. W. Incorporating social knowledge structures into computational models. Nature Communications 13, 6205 (2022).

13 Zhang, L., Lengersdorff, L., Mikus, N., Gläscher, J. & Lamm, C. Using reinforcement learning models in social neuroscience: frameworks, pitfalls and suggestions of best practices. Social Cognitive and Affective Neuroscience 15, 695–707 (2020).

14 Lockwood, P. L., Apps, M. A., Valton, V., Viding, E. & Roiser, J. P. Neurocomputational mechanisms of prosocial learning and links to empathy. Proceedings of the National Academy of Sciences 113, 9763–9768 (2016).

15 O’Connell, K., Walsh, M., Padgett, B., Connell, S. & Marsh, A. A. Modeling variation in empathic sensitivity using go/no-go social reinforcement learning. Affective Science 3, 603–615 (2022).

16 Zaki, J., Kallman, S., Wimmer, G. E., Ochsner, K. & Shohamy, D. Social cognition as reinforcement learning: feedback modulates emotion inference. Journal of Cognitive Neuroscience 28, 1270–1282 (2016).

17 Heffner, J., Son, J. Y. & FeldmanHall, O. Emotion prediction errors guide socially adaptive behaviour. Nature human behaviour 5, 1391–1401, doi:10.1038/s41562-021-01213-6 (2021).

18 Ruff, C. C. & Fehr, E. The neurobiology of rewards and values in social decision making. Nature Reviews Neuroscience 15, 549–562 (2014).

19 Arabadzhiyska, D. H. et al. A common neural currency account for social and non-social decisions. bioRxiv, 2021.2010. 2018.464762 (2021).

20 Olsson, A., Knapska, E. & Lindström, B. The neural and computational systems of social learning. Nature Reviews Neuroscience 21, 197–212 (2020).

21 Rogers-Carter, M. M. et al. Insular cortex mediates approach and avoidance responses to social affective stimuli. Nature neuroscience 21, 404–414 (2018).

22 Feng, C. et al. Common brain networks underlying human social interactions: Evidence from large-scale neuroimaging meta-analysis. Neuroscience & Biobehavioral Reviews 126, 289–303 (2021).

23 Gehrlach, D. A. et al. Aversive state processing in the posterior insular cortex. Nature neuroscience 22, 1424–1437 (2019).

24 Sanfey, A. G., Rilling, J. K., Aronson, J. A., Nystrom, L. E. & Cohen, J. D. The neural basis of economic decision-making in the ultimatum game. Science 300, 1755–1758 (2003).

25 Corlett, P. R., Mollick, J. A. & Kober, H. Meta-analysis of human prediction error for incentives, perception, cognition, and action. Neuropsychopharmacology 47, 1339–1349 (2022).

26 Behrens, T. E., Hunt, L. T. & Rushworth, M. F. The computation of social behavior. science 324, 1160–1164 (2009).

27 Apps, M. A., Rushworth, M. F. & Chang, S. W. The anterior cingulate gyrus and social cognition: tracking the motivation of others. Neuron 90, 692–707 (2016).

28 Lockwood, P. L. & Wittmann, M. K. Ventral anterior cingulate cortex and social decision-making. Neuroscience & Biobehavioral Reviews 92, 187–191 (2018).

29 Chang, S. W., Gariépy, J.-F. & Platt, M. L. Neuronal reference frames for social decisions in primate frontal cortex. Nature neuroscience 16, 243–250 (2013).

30 Gan, X. et al. A neurofunctional signature of subjective disgust generalizes to oral distaste and socio-moral contexts. Nature human behaviour, 1–20 (2024).

31 Zhou, F. et al. A distributed fMRI-based signature for the subjective experience of fear. Nature communications 12, 6643 (2021).

32 Kragel, P. A., Koban, L., Barrett, L. F. & Wager, T. D. Representation, pattern information, and brain signatures: from neurons to neuroimaging. Neuron 99, 257–273 (2018).

33 Woo, C.-W., Chang, L. J., Lindquist, M. A. & Wager, T. D. Building better biomarkers: brain models in translational neuroimaging. Nature neuroscience 20, 365–377 (2017).

34 Gabay, A. S., Radua, J., Kempton, M. J. & Mehta, M. A. The Ultimatum Game and the brain: A meta-analysis of neuroimaging studies. Neuroscience & Biobehavioral Reviews 47, 549–558 (2014).

35 Kohoutová, L. et al. Toward a unified framework for interpreting machine-learning models in neuroimaging. Nature protocols 15, 1399–1435 (2020).

36 Rangel, A., Camerer, C. & Montague, P. R. Neuroeconomics: The neurobiology of value-based decision-making. Nature Reviews. Neuroscience 9, 545 (2008).

37 Frömer, R., Dean Wolf, C. K. & Shenhav, A. Goal congruency dominates reward value in accounting for behavioral and neural correlates of value-based decision-making. Nature communications 10, 4926 (2019).

38 Kording, K. Decision theory: what “should” the nervous system do? Science 318, 606–610 (2007).

39 Kubanek, J. Optimal decision making and matching are tied through diminishing returns. Proceedings of the National Academy of Sciences 114, 8499–8504 (2017).

40 Kahneman, D. A psychological perspective on economics. American economic review 93, 162–168 (2003).

41 Heffner, J. & FeldmanHall, O. A probabilistic map of emotional experiences during competitive social interactions. Nature communications 13, 1718 (2022).

42 Villano, W. J., Otto, A. R., Ezie, C., Gillis, R. & Heller, A. S. Temporal dynamics of real-world emotion are more strongly linked to prediction error than outcome. Journal of Experimental Psychology: General 149, 1755 (2020).

43 Skvortsova, V., Palminteri, S. & Pessiglione, M. Learning to minimize efforts versus maximizing rewards: computational principles and neural correlates. Journal of Neuroscience 34, 15621–15630 (2014).

44 Kurniawan, I. T., Guitart-Masip, M., Dayan, P. & Dolan, R. J. Effort and valuation in the brain: the effects of anticipation and execution. Journal of Neuroscience 33, 6160–6169 (2013).

45 Hauser, T. U. et al. Temporally dissociable contributions of human medial prefrontal subregions to reward-guided learning. Journal of Neuroscience 35, 11209–11220 (2015).

46 Rushworth, M. F., Noonan, M. P., Boorman, E. D., Walton, M. E. & Behrens, T. E. Frontal cortex and reward-guided learning and decision-making. Neuron 70, 1054–1069 (2011).

47 Grant, R. I. et al. Specialized coding patterns among dorsomedial prefrontal neuronal ensembles predict conditioned reward seeking. Elife 10, e65764 (2021).

48 Corradi-Dell’Acqua, C., Tusche, A., Vuilleumier, P. & Singer, T. Cross-modal representations of first-hand and vicarious pain, disgust and fairness in insular and cingulate cortex. Nature communications 7, 10904 (2016).

49 Yao, S. et al. Oxytocin facilitates approach behavior to positive social stimuli via decreasing anterior insula activity. International Journal of Neuropsychopharmacology 21, 918–925 (2018).

50 Schaich Borg, J., Lieberman, D. & Kiehl, K. A. Infection, incest, and iniquity: Investigating the neural correlates of disgust and morality. Journal of cognitive neuroscience 20, 1529–1546 (2008).

51 Bryden, D. W., Johnson, E. E., Tobia, S. C., Kashtelyan, V. & Roesch, M. R. Attention for learning signals in anterior cingulate cortex. Journal of Neuroscience 31, 18266–18274 (2011).

52 Jezzini, A., Bromberg-Martin, E. S., Trambaiolli, L. R., Haber, S. N. & Monosov, I. E. A prefrontal network integrates preferences for advance information about uncertain rewards and punishments. Neuron 109, 2339–2352. e2335 (2021).

53 Kuehn, E., Mueller, K., Lohmann, G. & Schuetz-Bosbach, S. Interoceptive awareness changes the posterior insula functional connectivity profile. Brain Structure and Function 221, 1555–1571 (2016).

54 Greene, J. D., Nystrom, L. E., Engell, A. D., Darley, J. M. & Cohen, J. D. The neural bases of cognitive conflict and control in moral judgment. Neuron 44, 389–400 (2004).

55 Buckholtz, J. W. et al. The neural correlates of third-party punishment. Neuron 60, 930–940 (2008).

56 Civai, C., Miniussi, C. & Rumiati, R. I. Medial prefrontal cortex reacts to unfairness if this damages the self: a tDCS study. Social Cognitive and Affective Neuroscience 10, 1054–1060 (2015).

57 Zoh, Y., Chang, S. W. & Crockett, M. J. The prefrontal cortex and (uniquely) human cooperation: a comparative perspective. Neuropsychopharmacology 47, 119–133 (2022).

58 Gao, X. et al. Distinguishing neural correlates of context-dependent advantageous-and disadvantageous-inequity aversion. Proceedings of the National Academy of Sciences 115, E7680–E7689 (2018).

59 Bellucci, G., Feng, C., Camilleri, J., Eickhoff, S. B. & Krueger, F. The role of the anterior insula in social norm compliance and enforcement: Evidence from coordinate-based and functional connectivity meta-analyses. Neuroscience & Biobehavioral Reviews 92, 378–389 (2018).

60 Dupont, L., Santangelo, V., Azevedo, R. T., Panasiti, M. S. & Aglioti, S. M. Reputation risk during dishonest social decision-making modulates anterior insular and cingulate cortex activity and connectivity. Communications biology 6, 475 (2023).

61 Grecucci, A., Giorgetta, C., Van’t Wout, M., Bonini, N. & Sanfey, A. G. Reappraising the ultimatum: an fMRI study of emotion regulation and decision making. Cerebral cortex 23, 399–410 (2013).

62 Chowdhury, R. et al. Dopamine restores reward prediction errors in old age. Nature neuroscience 16, 648–653 (2013).

63 Martins, D., Lockwood, P., Cutler, J., Moran, R. & Paloyelis, Y. Oxytocin modulates neurocomputational mechanisms underlying prosocial reinforcement learning. Progress in neurobiology 213, 102253 (2022).

64 Xu, T. et al. The central renin–angiotensin system: A genetic pathway, functional decoding, and selective target engagement characterization in humans. Proceedings of the National Academy of Sciences 121, e2306936121 (2024).

65 Xu, T. et al. Angiotensin blockade enhances motivational reward learning via enhancing striatal prediction error signaling and frontostriatal communication. Molecular Psychiatry 28, 1692–1702 (2023).

66 Tay, R. Correlation, variance inflation and multicollinearity in regression model. Journal of the Eastern Asia Society for Transportation Studies 12, 2006–2015 (2017).

67 Li, J. et al. Common and dissociable contributions of alexithymia and autism to domain-specific interoceptive dysregulations: A dimensional neuroimaging approach. Psychotherapy and Psychosomatics 88, 187–189 (2019).

68 Xu, X. et al. Intrinsic connectivity of the prefrontal cortex and striato-limbic system respectively differentiate major depressive from generalized anxiety disorder. Neuropsychopharmacology 46, 791–798 (2021).

69 Davis, T. et al. What do differences between multi-voxel and univariate analysis mean? How subject-, voxel-, and trial-level variance impact fMRI analysis. Neuroimage 97, 271–283 (2014).

70 Slotnick, S. D. Cluster success: fMRI inferences for spatial extent have acceptable false-positive rates. Cognitive neuroscience 8, 150–155 (2017).

