## Supplemental Information for "Distinct neural computations scale the violation of expected reward and emotion in social transgressions"

Benjamin Becker

Department of Psychology, The University of Hong Kong, Hong Kong, China

### Supplemental Methods

#### MRI acquisition and preprocessing

MRI data were acquired on a 3.0-T GE Discovery MR system (General Electric Medical System, Milwaukee, WI, USA). The high-resolution brain anatomical MRI image were acquired using a T1-weighted sequence (Time repetition (TR)=6ms; echo time=2ms; flip angle=9°; field of view=256×256 mm; matrix size=256×256; voxel size=1×1×1 mm; number of slices, 156; slice thickness, 1 mm) to improve spatial normalization of the functional data and exclude subjects with apparent brain pathologies. Functional data using blood oxygenation level-dependent contrast were obtained on this scanner using a T2*-weighted echo planar imaging sequence (TR=2000ms; echo time=30ms; flip angle=90°; field of view=240×240 mm; voxel size=3.75×3.75×3 mm; number of slices, 39; slice thickness, 3 mm; resolution=64×64).

All fMRI images were preprocessed and analyzed using standard workflows in SPM 12 (Statistical Parametric Mapping; http://www.fil.ion.ucl.ac.uk/spm/; Welcome Trust Centre for Neuroimaging). The first 5 volumes of each functional time series were discarded to allow for T1 equilibration. Remaining images were corrected for acquisition time delay, realigned to correct for head motion, unwarped for magnetic field inhomogeneities correction, and co-registered with the T1-weighted structural image. After that the images were normalized to Montreal Neurological Institute standard space (interpolated to 2×2×2 mm voxel size) using the segmentation parameters from the anatomical images, and were then spatially smoothed using an isotropic Gaussian kernel with full-width at half-maximum of 8 mm.

#### Brain predictive patterns of reward and emotional PEs

With the purpose of obtaining sensitive neural patterns of brain activity that predict reward and emotional PEs, we employed whole-brain multivariate machine-learning pattern analyses. Specifically, we sorted individual trial-based reward and emotional PE values with descend sequence and divided them into 5 levels (i.e., 5, 4, 3, 2, 1). In the general liner models (GLM) for reward and emotional PE, separately, we included five separate regressors time-logged to the presentations of actual reward offers (or actual emotional ratings) for each reward PE level (or emotional PE level) (i.e., 1-5), to model brain activity in response to each reward or emotional PE level separately. The six estimated head motion parameters were added as nuisance variable in the GLM models.  We then utilized a support vector regression algorithm (linear kernel with C= 1) implemented in the Spider toolbox (http:// people.kyb.tuebingen.mpg.de/spider) with individual beta maps (one per PE level for each subject) as features to predict participants true PE values. To evaluate the performance of the predictive pattern of reward and emotional PEs (trained on the whole brain), we assessed overall (between- and within-subjects; 43×5=215 pairs in total) and within-subject (5 pairs per subject) Pearson correlations (r) between the cross-validated predictions and the actual ratings to evaluate the effect sizes. Moreover, we computed the classification accuracies of the PE predictive patterns between low, moderate and high PE levels (i.e., high versus low, high versus moderate, and moderate versus low) via the use of forced-choice classification, where signature responses were compared for two conditions tested within the same participant (the higher indicated large violations of predicted emotions or rewards). Two-sided binomial tests were used to test whether the classification accuracies were higher than chance-level (50%).

### Supplemental Results

| Variable | Estimate(s.e) | z | p |
|---|---|---|---|
| Punitive decision | | | |
| Intercept | 0.17(0.66) | 0.18 | 0.86 |
| Gender | 0.94(0.89) | 1.07 | 0.29 |
| Unfairness | 4.06(0.52) | 7.84 | **<0.001** |
| Gender*Unfairness | 0.03(0.64) | 0.05 | 0.96 |

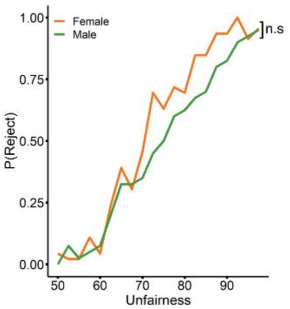


**Fig. S1** Unfairness predicts the punishment decisions regardless of gender. n.s – not significant

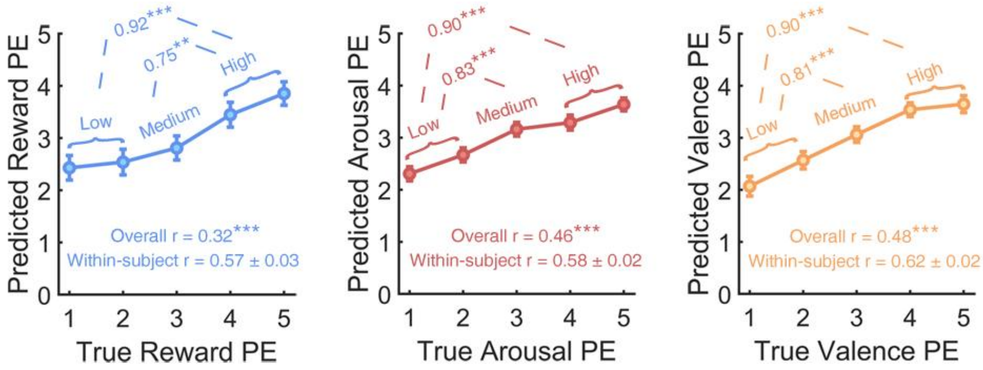


**Fig. S2** Multivariate neural pattern predicted PEs compared to actual PEs. Accuracy provided for forced-choice comparisons. P values based on two-sided independent binomial tests. r indicates Pearson correlation coefficient between predicted and true reward or emotional PEs. ***p<0.001, **p<0.01

**Table S1**. Brain regions for reward prediction error under accept decisions.

| Regions | x | y | z | k | $P_{FDR}$ |
|---|---|---|---|---|---|
| Ventromedial prefrontal cortex | 0 | 32 | -8 | 66 | <0.05 |
| Dorsolateral prefrontal cortex | -42 | 36 | 30 | 352 | <0.05 |
| Dorsomedial prefrontal cortex | -4 | 52 | 38 | 115 | <0.05 |
